# Detection of a Unique Unfolded p53 Conformation (U-p53^AZ^) in the Tissue of Alzheimer’s Disease Patients

**DOI:** 10.1101/2025.08.05.668780

**Authors:** David Lynch, Sam Agus, Rakez Kayed, Michael Rasche, Madison Samples, Simona Piccirella

## Abstract

**INTRODUCTION:** Alzheimer’s Disease (AD) is a major public health challenge. An AD-specific unfolded form of the p53 protein was identified (U-p53^AZ^) and data from animal studies suggests it colocalizes with pTau, in neurofibrillary tangles (NFT)

**METHODS:** Brain samples of patients with AD, cognitively unimpaired non-AD, and with fronto-temporal dementia (FTD) are being tested, comparing the pattern of U-p53^AZ^ accumulation (using the antibody 2D3A8, developed by Diadem SpA) with accumulation of pTau, beta-amyloid, and TDP-43.

**RESULTS:** U-p53^AZ^ accumulates in brain samples from AD, colocalizes with pTau in the NFT, with a small degree of accumulation in neuritic plaques (NP). The intensity of staining correlates with the pTau staining and Braak staging. No staining is observed in samples from cognitively unimpaired non-AD and FTD.

**DISCUSSION:** This study provides further support to the hypothesis that U-p53^AZ^ is a player in AD pathology, in link with pTau accumulation and Braak staging.

## Introduction

Alzheimer’s disease (AD) is a devastating neurodegenerative disorder that will affect approximately 107 million patients by 2050 [1]. This will have a huge cost, estimated to be 16.9 trillion US dollars combined across all countries [2]. The development of novel treatments has given some hope that this disease can be controlled [3]. Because these treatments are approved for patients with early symptomatic AD (i.e. minimal cognitive impairment (MCI) or mild dementia, due to AD), and because of other limitations that relate to the safety profile of these treatments, only about 1 million individuals are, currently, treatment eligible in the United States [4, 5].

With more than half of the population above the age of 65 expressing concerns about their cognitive health [6], there is a need for new biomarkers which can help in management of these individuals, setting priorities (when the tests read as positive), and providing peace of mind (when they read negative). These biomarkers should have a good prognostic correlation (i.e. determining the risk for clinically significant deterioration and providing the potential timeframe for this deterioration), be able to be tested with minimal invasiveness (as opposed to CSF testing), be easily accessible (as opposed to PET scans and CSF testing), and are not excessively expensive.

Buizza et al [7] showed that under AD conditions, p53 undergoes a conformational change and Abate et al described a specific conformational variant as clinically relevant in AD progression and its role as a biomarker in the AD diagnosis[8]. An AD-specific form of the unfolded p53 protein (U-p53^AZ^) has been identified [7-9]. An antibody to U-p53^AZ^ (2D3A8) was developed, to detect this variant in the plasma of individuals with AD and performs well in predicting the risk for a clinically significant deterioration within the next 6 years in both Cognitive unimpaired and Mild Cognitive Impaired patients at baseline [10]. As can be seen in Figure 1, the unfolding of the p53 protein permits the access of the antibody to the binding site in the protein.

**Figure 1:**
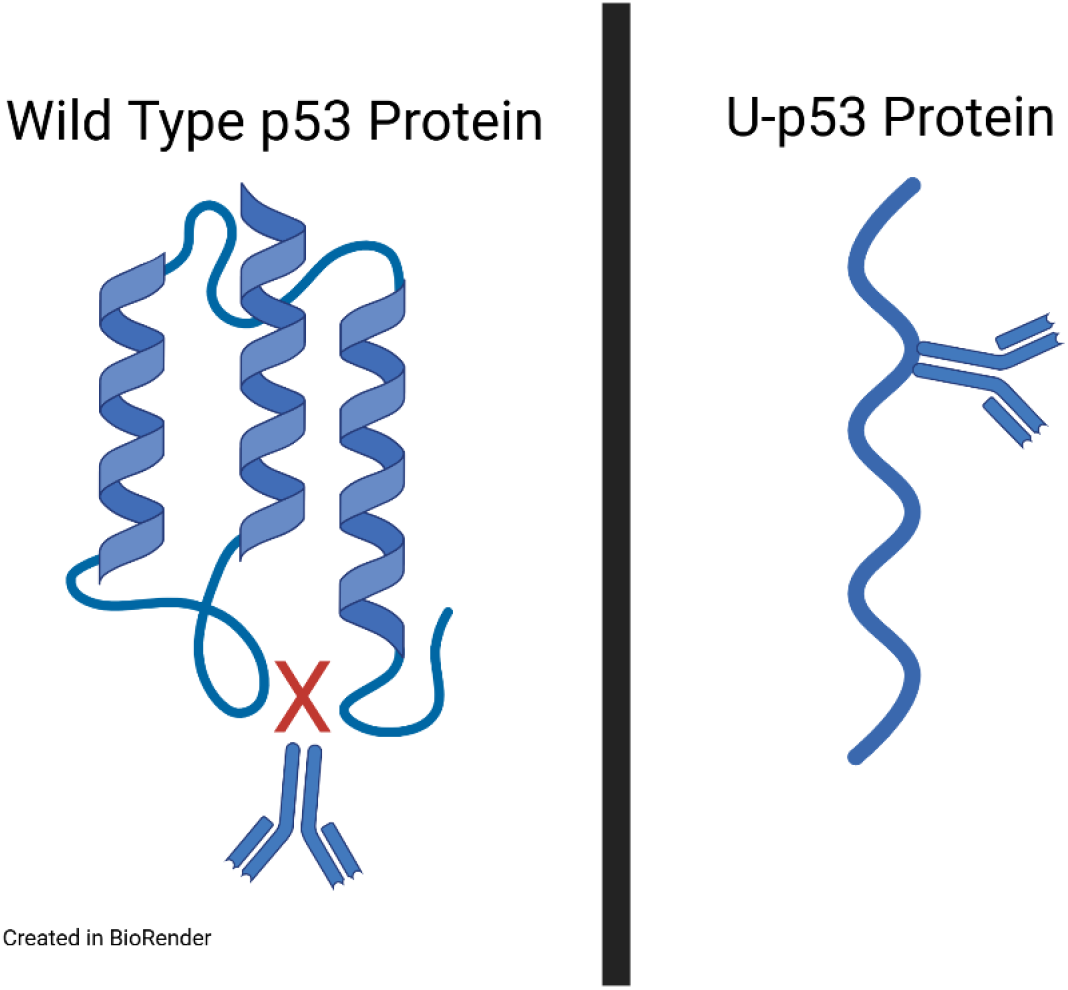
Comparison of U-p53^AZ^ Antibody Binding in Wild Type and Unfolded p53 Proteins.

The p53 protein is well-known to be associated with neurodegenerative diseases [11]. Under normal circumstances, p53 exists as a tetrameric protein that serves to, amongst other functions, repair DNA damage and regulate cellular senescence [12, 13]. The protein is, itself, inherently disordered, meaning that it easily forms alternative structures thanks to the composition of its amino acids [14]. The loss of p53’s normal structure allows it to quickly form alternatives, and it is likely that these have clinical implications [15, 16]. Farmer et al found phosphorylated p53 and phosphorylated tau, a known contributor to AD, to be co-localized, suggesting that not only is p53 active in AD, it also directly interacts with one of the main known drivers of AD. As the tau protein is known to interact with other proteins, forming hetero-aggregates [17], and for disease-associated forms to spread in a prion-like manner, it is possible that U-p53^AZ^ is formed in response to the changing environment caused by the development of AD, triggered by the development of tau aggregates.

Thus far, however, the unique protein conformation of U-p53^AZ^ has not been examined in human brain tissue from AD patients. The demonstration of this protein being present in tissue from patients with diagnosed AD would strengthen the association between U-p53^AZ^ and AD, demonstrating that it is present in brain tissue as well as peripheral blood of AD patients. This would not only strengthen the utility of the U-p53^AZ^ antibody as a prognostic biomarker, it would also further demonstrate the integral role that p53 plays in the development of AD.

The aim of this study is exploring the expression of U-p53^AZ^ in the brain from AD patients. Furthermore, due to its role as prognostic biomarker of AD progression, we have investigated its correlation with the severity of AD increases.

## Materials and Methods

### Manufacture of Recombinant p53 Material

Recombinant full-length p53 wild type protein, (WT-p53), MW 43.7 kDa, was expressed in an *E. coli* BL21(DE3) system (New England Biolabs, Cat#C2527). The protein was purified by using sonication to lyse the cells in lysis buffer (made with molecular grade water (Corning, Cat.# 46-000-CM) to dilute to1 M NaCl (Corning, Cat.# 46-032-CV), 25 mM Tris.HCl (Corning, Cat.# 46-031-CM)) in the presence of protease inhibitor (Roche, Cat. #11873580001) and phenylmethanesulfonyl fluoride (PMSF) (Sigma-Aldrich, Cat.#P7626). Cell debris was removed by centrifugation. Unwanted proteins were removed by the use of a NiSO_4_ resin (His-Pur Ni-NTA Resin, Thermo Scientific, Cat.# 88223) column to bind the protein. The His-tag of the protein was removed via on-column digestion with thrombin (Millipore, Cat.# 69671-3) for 48 hours at 4° C. The captured protein was then purified with fast protein liquid chromatography (Amersham Biosciences, Explorer 10) using a size-exclusion chromatography column (Superdex 75 Increase 10/300 GL, Cytiva life sciences Cat.# 29148721). Protein samples were stored at -20° C. in 1x phosphate buffered saline (PBS) (Corning, Cat.# 46-013-CM) with 0.02% sodium azide (Acros, Cat.# UN1687). The concentration of the protein samples was confirmed using a Pierce’s bicinchoninic acid (BCA) assay (ThermoFisher, Cat.# 23235).

### Preparation of Unfolded p53 Protein (U-p53)

The unfolded p53 (U-p53) was prepared similarly to the process described by Butler & Loh (2003)[18] by adding previously prepared p53 protein to 10% acetic acid (Acros, Cat.# 42322-0025) and 0.5 M EDTA (Fisher, Cat. # 15575020) in a roughly 96:3:1 ratio. Immediately, the solution was placed on ice and incubated for roughly 1 minute before 0.5 M Tris.HCl, made by diluting 1 M Tris.HCl in water, was added to the solution, at a volume of 1.5x the initial volume of the solution. The solution was then desalted using a PD10 column (Cytiva, Cat. #GE17-0851-01). Solution was then stored in 10% glycerol (Sigma, Cat. #G5516-1L) and 0.02% sodium azide solution in 1X PBS solution. U-p53 obtained following this protocol is used as surrogate of the clinical U-p53^AZ^.

### Western blots

1.0 µg of the U-p53 and WT-p53, and precision protein dual color standard (Bio-Rad, Cat.# 1610374) to serve as a protein ladder, were loaded in two separate groups and resolved on a precast NuPAGE 4-12% Bis-Tris gel for SDS-PAGE (Invitrogen, Cat.# NP0335 BOX) with an XCell SureLock Mini gel tank (Invitrogen, Cat.# EI0001) and transferred to nitrocellulose membranes (Bio-Rad, Cat.# 1620115) with the use of a mini trans-blot cell (Bio-Rad, Cat.# 170390). Membranes were blocked with 10% nonfat milk (BioKEMIX, Cat.# M-0842) in 1x Tris-buffered saline solution (Corning, Cat.# 46-012-CM) plus 0.1% Tween-20 (Fisher, Cat.# BP337-500) (TBST) for 1.5 hours at room temperature. The membranes were then divided so that each half of the membrane contained one lane with a protein ladder, U-p53, and WT-p53. The membranes were then incubated overnight at 4° C., one membrane half with anti-p53 antibody DO-7 (Cell Signaling, Cat.# 48818) at a 1/1000 concentration and the other with anti-U-p53^AZ^ antibody 2D3A8 (provided by Diadem SpA) at a concentration of 0.5 µg/mL. Membranes were then incubated with, respectively, horseradish peroxidase goat anti-mouse IgG secondary antibody (Bio-Rad, Cat.# 170-6516) and goat anti-mouse IgM secondary antibody (Jackson ImmunoResearch Laboratories, Cat.# 115-001-020). WesternBright ECL (Advansta, Cat.# 12045-D50) was used to detect the signal.

### Human AD samples

Human AD samples are listed in Table 1. Braak staging was performed by clinical pathologists. Samples were fixed in optimal cutting temperature (OCT) compound (Tissue-Tek, Cat.# 4583) and subsequently sectioned on a CryoStat (Epredia, CryoStar NX50). Sections were stored at -80° C. until use.

**Table 1:**
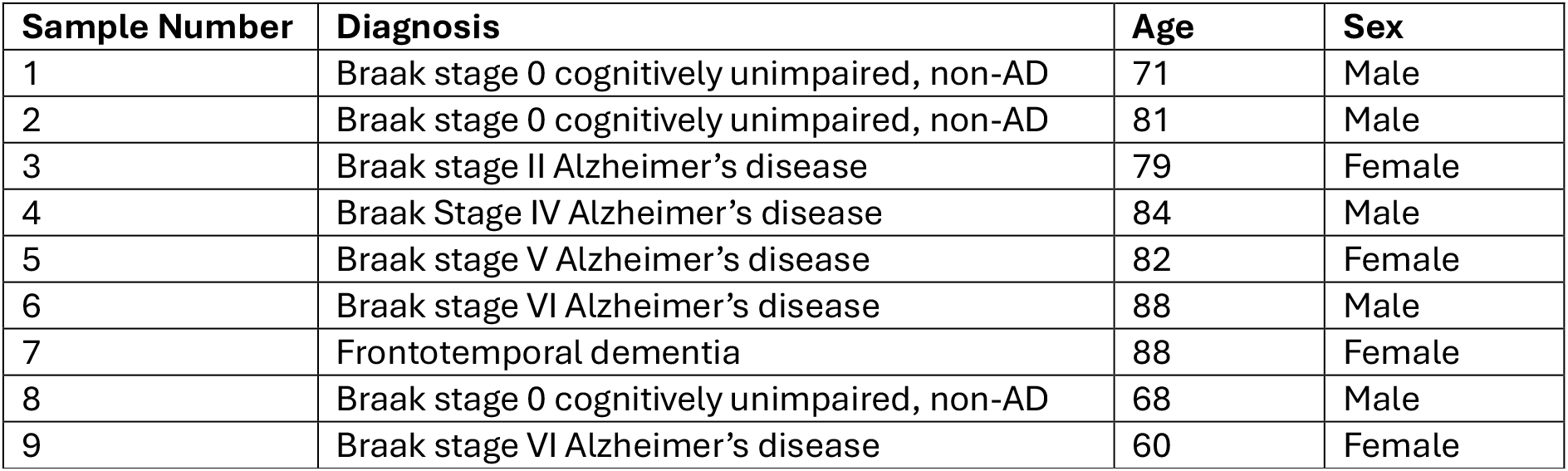
Patient Sample List.

| Sample Number | Diagnosis | Age | Sex |
| --- | --- | --- | --- |
| 1 | Braak stage 0 cognitively unimpaired, non-AD | 71 | Male |
| 2 | Braak stage 0 cognitively unimpaired, non-AD | 81 | Male |
| 3 | Braak stage II Alzheimer's disease | 79 | Female |
| 4 | Braak Stage IV Alzheimer's disease | 84 | Male |
| 5 | Braak stage V Alzheimer's disease | 82 | Female |
| 6 | Braak stage VI Alzheimer's disease | 88 | Male |
| 7 | Frontotemporal dementia | 88 | Female |
| 8 | Braak stage 0 cognitively unimpaired, non-AD | 68 | Male |
| 9 | Braak stage VI Alzheimer's disease | 60 | Female |

### AD Patients-Immunofluorescence Staining

Slides from AD patients listed in Table 1 were stained for the co-presence of a variety of cellular markers, including U-p53^AZ^, phosphorylated tau protein, phosphorylated p53 protein, and for the presence of beta sheets (often found in toxic oligomers). Antibody concentrations are shown in Table 2. The slides were first fixed in 100% chilled methanol (Fisher, Cat.# 61009-0040), washed in 1X PBS (pH 7.4) and then the tissue was circled with a PAP pen (Ihc World LLC, Cat.# NC0552848). The slides were then blocked in 5% goat serum (Sigma, Cat.# G9023-10ML) diluted in PBS with 0.25% Triton X-100 (Sigma Aldrich, Cat.# T8787-100ML). The slides were washed with 70% ethanol (purified water was used to dilute ethyl alcohol absolute, Sigma, Cat.# 459844), and then 1x TrueBlack lipofuscin autofluorescence quencher (Biotium, Cat.# 23007) was applied to each slide. Slides were washed with ethanol until the ethanol ran clear, then the slides were washed with 1X PBS. Slides were then stained with the first primary antibody, 2D3A8 (specific to detect U-p53^AZ^), overnight at 4°C., and then washed with PBS. The slides were then incubated with secondary antibody, goat anti mouse IgM (Licor, Cat.# 926-32280), for 1 hour at room temperature. Samples were then blocked with F(abs) (Jackson Immunoresearch, Cat.# 715-007-003) and then the slides were incubated for 1 hour at 37° C. with two other primary antibodies, after ensuring that the two isotypes could be used simultaneously. The slides had 1X TrueBlack lipofuscin autofluorescence quencher applied to it again. The slides were washed in 70% ethanol, then washed in PBS. The slides were then stained with 1mg/mL DAPI (Invitrogen, Cat.# D3571) diluted at 1/1000 in PBS. Slides were then washed in PBS and then mounted with ProLong Gold Antifade mounting media (ThermoFisher, Cat.# P10144) and sealed with nail polish.

**Table 2:**
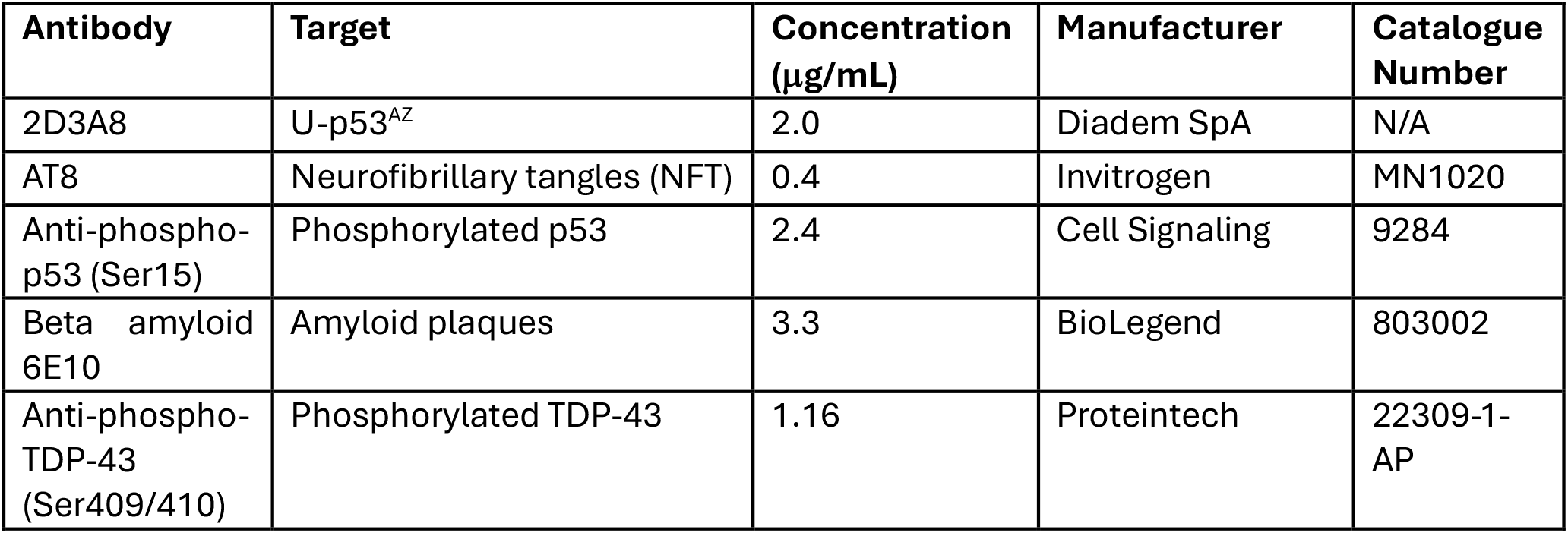
List of Antibodies and Targets in Immunofluorescence. The 2D3A8 antibody is not currently available commercially. It is manufactured and developed by Diadem SpA. as a prognostic biomarker, to be utilized for individuals with cognitive concerns or MCI as part of the AlzoSure® Predict kit.

### Antibody Selection

The antibodies used are outlined in Table 2. Antibodies were selected that can detect specific disease-associated conformations or phosphorylated states of p53, Tau, TDP-43 and amyloid beta proteins.

### Statistical Analysis

The proportion of the Western blot image that is positive both for DO-7 and for 2D3A8 was calculated with the use of the FIJI plug-in for the ImageJ program.

## Results

### Western Blot of WT-p53 and U-p53

The Western blot of the U-p53^AZ^ control protein shows that 2D3A8 does bind preferentially to the unfolded p53 protein as opposed to normal, p53 WT tetramer. The p53 DO7 antibody is used to confirm the presence of p53 in its WT sequence. Figure 2 shows representative Western blots for the process, clearly indicating that there is U-p53^AZ^ expression, but not to the WT. Analysis on ImageJ confirms that there is no staining of the WT p53 by 2D3A8 Ab in our control proteins. The implications of this are that the 2D3A8 antibody will specifically bind to the unfolded p53 protein. While signal detection is limited compared to that detected by DO-7, this does still indicate that there is differentiation between the two conformations of the protein.

**Figure 2:**
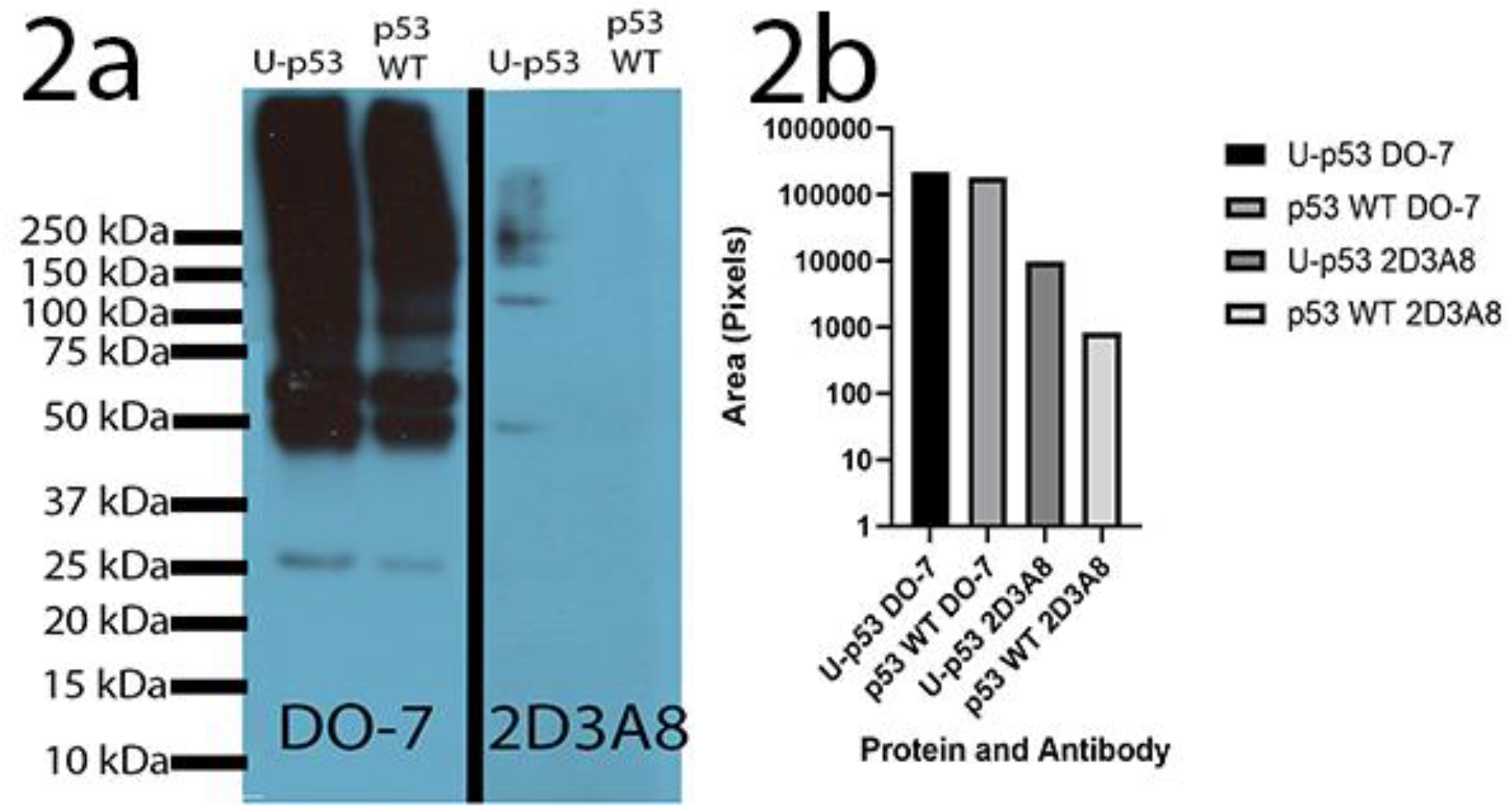
Western blot and analysis comparing signal detection of U-p compared to WT-p53 using the 2D3A8 antibody.

### Immunofluorescence in samples from patients with AD and cognitively unimpaired, non-AD controls

Figure 3 shows staining results from samples of patients with AD, compared to samples from a healthy control. Samples from AD patients show, as expected, an increase in the level of pTau staining intensity (see panels 3Id, 3IIId, and 3IVd), in correlation with the Braak stage. Staining with the 2D3A8 antibody shows the U-p53^AZ^ colocalizes with pTau, in the cell bodies, which in line with previous work in AD animal models.

**Figure 3:**
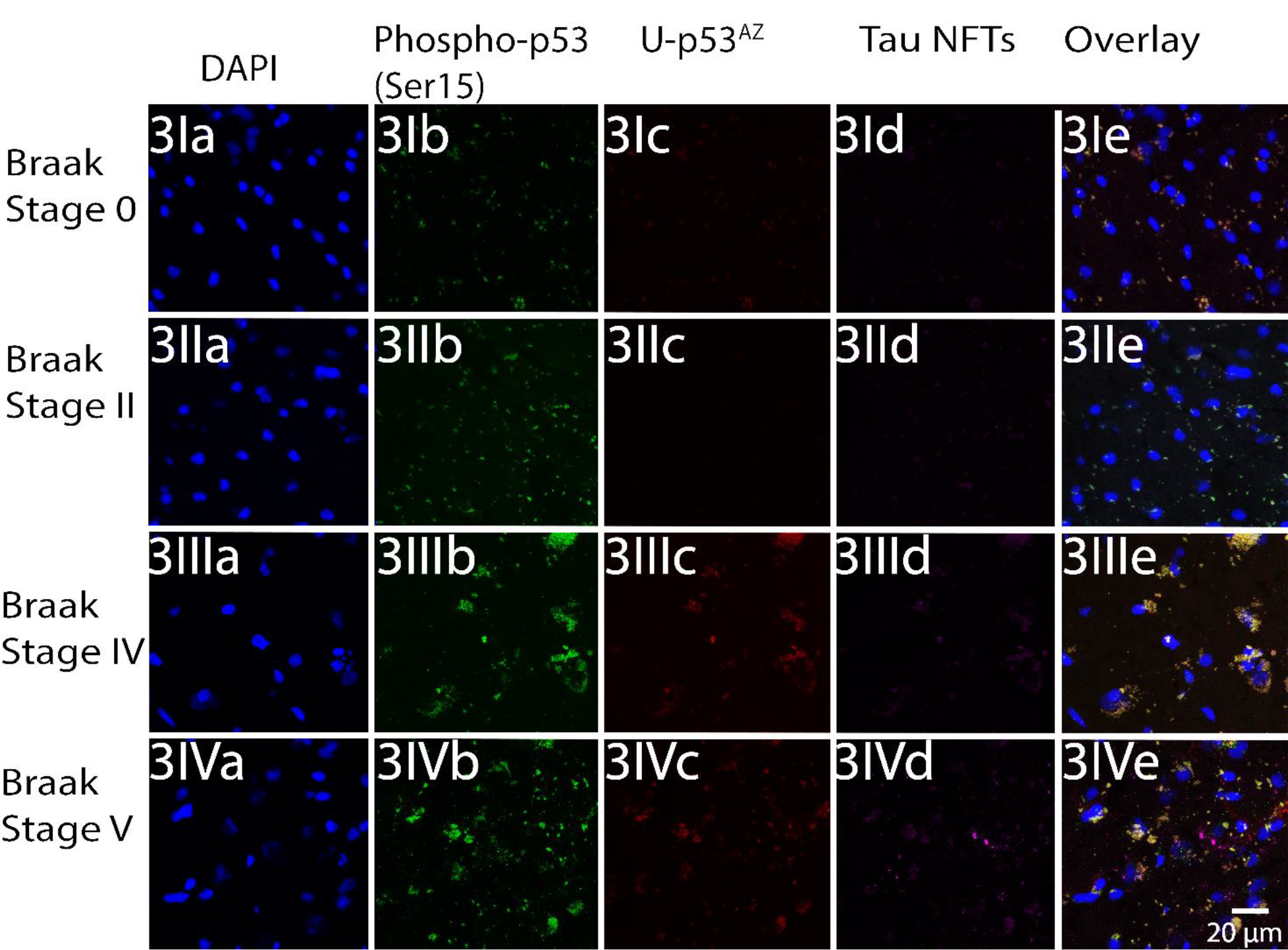
Immunofluorescence in samples from Alzheimer’s Disease (AD) Patients Frozen Sections. Figure 3I-3IV shows immunofluorescent staining from AD patients. In order, these patients were Braak stage 0, II, IV and V. Images marked ‘a’ show staining with DAPI to indicate the nuclei of the cells. Images marked ‘b’ show staining with the anti-phosphorylated p53 (Ser15). Images marked ‘c’ show staining with the anti-U-p53^AZ^ antibody 2D3A8. Images marked ‘d’ show staining with the anti-tau neurofibrillary tangle antibody AT8. Images marked ‘e’ show the integrated images of all four antibody stains.

Staining intensity of U-p53^AZ^ increases with the Braak staging and the intensity of pTau staining. There is no staining of U-p53^AZ^ in samples from the cognitively unimpaired non-AD control.

It is important to note that, at Braak Stage VI – representing the most advanced stage of AD pathology, the signal intensity of pTau and with the 2D3A8 antibody, while significantly higher than what is seen in samples from healthy controls, is also somewhat lower than the signal seen (for both targets), in Braak stage IV.

In addition, while the signal with the 2D3A8 antibody, colocalizes in the cell bodies, with the pTau signal – indicating that U-p53^AZ^ is present in the NFT, some signal can be also seen outside of the neurons, in colocalization with beta-amyloid staining (Figure 4), indicating that some of the unfolded p53 also accumulates in the neurotic plaques. This phenomenon, of minor accumulation outsides of the neurons, is not seen in samples from healthy controls, whereby there is also no beta-amyloid staining.

**Figure 4:**
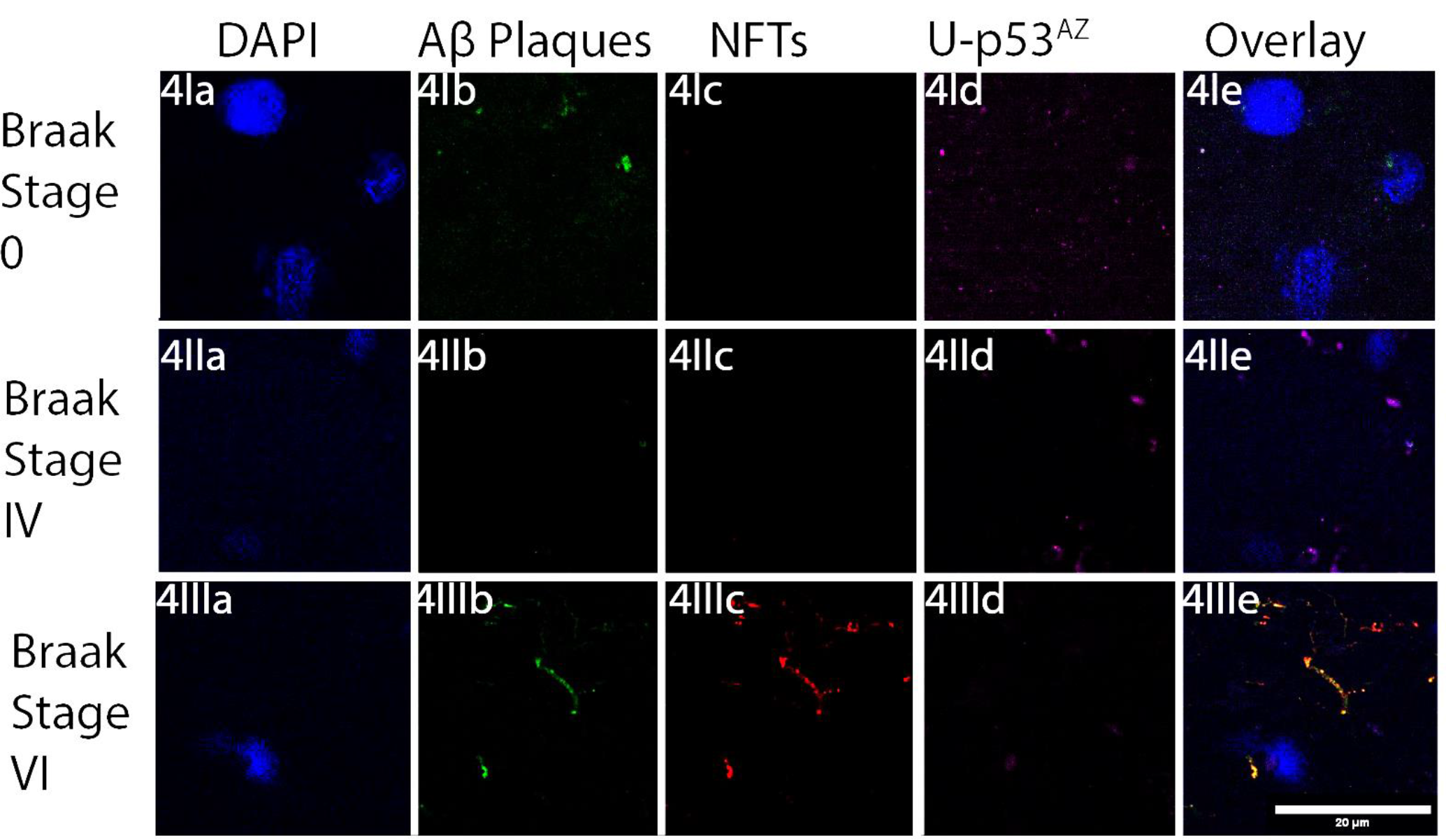
Immunofluorescent Staining of AD Patients from Braak Stages 0, IV and VI showing amyloid beta plaques (all panels marked ‘b’), neurofibrillary tangles (all panels marked ‘c) and U-p53^AZ^ (all panels marked ‘d’) staining. DAPI staining was shown in panels marked ‘a’. Panels marked ‘e’ show the overlay of the DAPI, Aβ plaque, neurofibrillary tangle and U-p53^AZ^ staining. Owing to a variable degree of background staining, the filter for the DAPI images was adjusted between each Braak stage patient.

### Immunofluorescence in samples from patients with fronto-temporal dementia (FTD)

Figure 5 compares the staining of the frontotemporal dementia (FTD) marker of phosphorylated TDP-43 (Ser409/410) from patients with AD and FTD and from a healthy control, comparing this to the staining for U-p53^AZ^ in the same slides. The AD sample comes from an individual with Braak stage VI disease. Samples were stained with an antibody for TDP-43 phosphorylated at Serine 409/410, a marker for FTD that is also known to stain positively in AD cases, and with the 2D3A8 antibody (to assess if U-p53^AZ^ is expressed also in FTD patients). While no signal is detected for phospho-TDP-43 and Up53^AZ^ in the healthy control sample, they are both detected in the AD sample. In the FTD sample, while some phospho-TDP-43 staining can be detected, there is no signal with the 2D3A8 antibody, indicating lack of accumulation of U-p53^AZ^.

**Figure 5:**
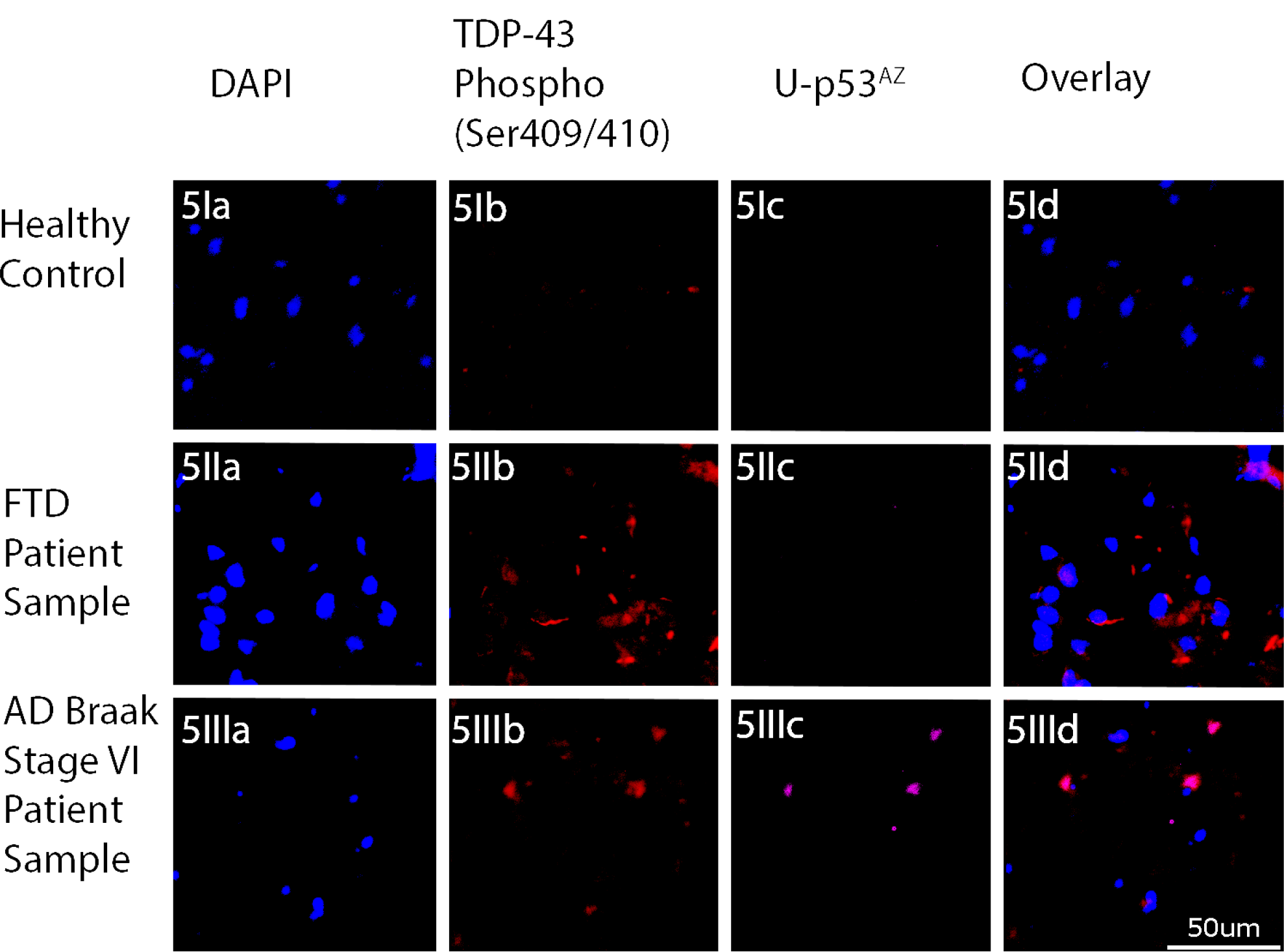
AD and FTD staining; panels with ‘b’ shows TDP-43 (Ser 409/410 phosphorylated) staining. Panels with ‘c’ show U-p53^AZ^ staining

## Discussion

It has been already demonstrated that U-p53^AZ^, specifically detected by 2D3A8 Ab, in blood it is able to distinguish between patients with and without AD [8, 9]. The significance of this study has been to show that there is a clear link between elements of AD pathology, specifically pTau.

Accumulation of U-p53^AZ^, as demonstrated by fluorescence staining with the proprietary antibody 2D3A8 occurs in AD, and in proximity to accumulation of pTau in NFT. While some unfolded protein can also be found in neuritic plaques (NP), but the extent of accumulation is closely correlating with the extent of pTau accumulation and the Braak stage. This observation may be in line with the utility of this marker as prognostic.

The observation that there seems to be a decrease in both pTau and U-p53^AZ^ staining, in samples from individuals with Braak VI stage can reflect the increased degree of neuronal loss, due to advanced neurodegeneration. The observation of a decrease in pTau staining in this study is somewhat in contrast to earlier observations and requires further assessment. However, the intensity of staining in this study at Braak VI stage, remain significantly increased when compared to healthy controls. A few studies have shown that in advanced Braak stages, neuronal loss and a phenotypic form of NFT related to advanced AD (also known as ‘ghost’ NFT), could lead to a plateau in increase of staining intensity in Braak stages V-VI [19].

The observation that 2D3A8, does not stain in samples from FTD, could be in-line with the possibility that U-p53^AZ^ is specific for AD.

Overall, this work adds another set of evidence, to inform about the link of U-p53^AZ^ to AD pathology. While the actual role of this variant in the pathology of AD remains to be confirmed, it seems that U-p53^AZ^ is accumulating, mostly in NFTs (although a small amount also accumulates in NP) and in a correlation with the accumulation of pTau. The diagnostic guidelines view pTau as a staging biomarker[20], and this correlation provides an insight to the role of U-p53^AZ^ as a biomarker for the progression of AD.

Data from analysis of plasma of individuals with dementia suggests that U-p53^AZ^ is specific to AD and in this work, the lack of staining in FTD samples, supports this observation as well.

Further analysis of the performance of U-p53^AZ^, in individuals which are cognitively unimpaired, as well as in individuals with MCI due to AD, is ongoing.

## Informed Consent Statement

All samples were frozen and previously collected. All the ethical approval of different studies are reported in method section.

## Acknowledgment

We thank Diadem SpA for the access and the use of its own antibody 2D3A8 (Patent number 10,183,990).

We thank Dr Yu-Hsiu Wang of the Department of Biochemistry and Molecular Biology at UTMB for his assistance with the confocal microscopy.

This work was funded by the National Institute of Health grants: AG054025, AG10448132, AG072458 and the Mitchell Center for Neurodegenerative Diseases.

